# *Mycobacterium ulcerans* not detected by PCR on human skin in Buruli ulcer endemic areas of south eastern Australia

**DOI:** 10.1101/2023.03.29.534673

**Authors:** Anita Velink, Jessica L. Porter, Timothy P Stinear, Paul D. R. Johnson

**Author notes:** **Corresponding author:** Paul D. R. Johnson.

## Abstract

**Introduction:** *Mycobacterium ulcerans* (MU) causes Buruli ulcer (BU), a geographically restricted infection that can result in skin loss, contracture, and permanent scarring. Lesion-location maps compiled from more than 640 cases in south eastern Australia suggest biting insects are likely involved in transmission, but it is unclear whether MU is brought by insects to their target or if MU is already on the skin and inoculation is an opportunistic event that need not be insect dependent.

**Methods:** We validated a PCR swab detection assay and defined its dynamic range using laboratory cultured *M. ulcerans* and fresh pigskin. We invited volunteers in Buruli-endemic and non-endemic areas to sample their skin surfaces with self-collected skin swabs tested by IS2404 quantitative PCR.

**Results:** Pigskin validation experiments established a limit-of-detection of 0.06 CFU/cm^2^ at a qPCR cycle threshold (Ct) of 35. Fifty-seven volunteers returned their self-collected kits of 4 swabs (bilateral ankles, calves, wrists, forearms), 10 from control areas and 47 from endemic areas. Collection was timed to coincide with the known peak-transmission period of Buruli. All swabs from human volunteers tested negative (Ct ≥35).

**Conclusions:** *M. ulcerans* was not detected on the skin of humans from highly BU endemic areas.

**Author Summary:** Buruli ulcer (BU) incidence is increasing in temperate south eastern Australia. We have yet to develop public health programs to assist people avoid BU partly because the precise mode of transmission is contentious. Recent research has shown that environmental contamination with *M. ulcerans* (the cause of BU) is widespread in endemic areas as a result of faecal shedding from infected possums that live close to humans, although direct human-possum contact is rare. We investigated the possibility that the skin of humans in endemic areas could become transiently contaminated with *M. ulcerans* while outdoors. If this were the case then BU prevention programs could be developed around skin protection and regular washing/showering. To study this possibility, we developed a sensitive skin swab PCR-assay that we tested using a pigskin laboratory model so we could be confident in our results. We asked volunteers to collect their own skin swabs after spending at least 4 hours outside during the known period of peak Buruli transmission. Fifty seven volunteers returned swab sets for testing. Our results were negative. We did not find evidence that humans in our endemic zone have *M. ulcerans* contamination on their skin.

## Introduction

Buruli ulcer (BU) is a geographically restricted infection caused by *Mycobacterium ulcerans* [1]. Listed by WHO as a Neglected Tropical Disease, BU occurs in 33 countries but characteristically only in specific locations. BU is not a fatal condition but can cause severe tissue destruction if not diagnosed and managed effectively. Recent advances in treatment with antibiotics have improved the outlook for sufferers in Buruli-active zones [2-7] which currently include west and sub-Saharan Africa [8], tropical Northern Australia [9] and coastal and urban zones of temperate south eastern Australia [10].

Over the last 15 years we have shown that possums (tree-dwelling marsupials) are environmental reservoirs and amplifiers of MU in our local endemic areas [11, 12] and that mosquitoes trapped in these areas test positive by PCR at a rate of at least 4 per 1000 [13]. Calculated MU cell load in these mosquitoes is in the order of 100 cells per positive mosquito [13]. In laboratory experiments we have established that very small inoculae of *M. ulcerans* initiate infections in mice (as few as 3 CFU) [14]. In contrast, in an abraded-skin hairless guinea pig model direct contact of even high concentrations of *M. ulcerans* cells is not sufficient to establish an infection [15]. We have also reported that although new BU cases are most often diagnosed in Victoria in winter and spring, peak transmission occurs in summer and autumn [10]. The paradox of a summer-transmitted disease appearing in winter is explained by a long incubation period with a median of 4.5 months plus additional time taken to establish the diagnosis after a lesion first appears [16-18].

By aggregating more than 600 cases of Buruli to a single human body map we recognized that lesion location is non-random and matches parts of the body bitten by mosquitoes rather than areas that come into direct mechanical contact with the environment, although there is some overlap [19]. We have also investigated skin temperature variation as an explanation for this non-random distribution of Buruli lesions but found only a weak association with thermographically measured skin temperature [20].

As environmental contamination with MU-positive excreta is extensive in endemic areas [11, 12] it is conceivable that direct contamination from the environment is the primary step in transmission of MU to humans and that secondary inoculation occurs from insect bites or other penetrating trauma. In this study we have investigated the hypothesis that people living in highly BU-endemic areas in south eastern Australia have environmentally acquired *M. ulcerans* colonization/contamination on their skin as a first step which could then be followed by a range of chance inoculating events including but not restricted to biting insects.

## Methods

### Method Validation - bacterial isolate and culture conditions

*M. ulcerans* clinical isolate JKD8049 obtained from a patient in Victoria, Australia in 2004 was grown at 30°C in 7H9 Middlebrook broth supplemented with OADC (Becton Dickinson, Sparks, MD, USA) [21]. To establish bacterial concentration in the culture preparations used for the pigskin validation model, colony forming units (CFU) were calculated by spot plating 3μl volumes of serial 10-fold dilutions (10^−1^ to 10^−4^) of a JKD8049 culture onto Middlebrook 7H10 agar plates with a 5×5 grid marked. The spots were allowed to dry, the plates loosely wrapped in plastic bags and then incubated as above for 10 weeks before counting colonies. Data analysis was performed using GraphPad Prism v9.5.0.

### DNA extraction and quantitative PCR

DNA was extracted from pigskin and human skin swabs using a DNeasy PowerSoil kit (Qiagen). Procedural extraction control blanks (swabs with sterile water) were included to monitor potential PCR contamination in addition to no-template negative PCR controls. IS2404 quantitative PCR (qPCR) was performed using technical triplicates as described previously [22].

### Pigskin validation model and swabbing technique

Pigskin was purchased from a butcher at the Victoria Market (Melbourne, Australia). The pigskin was marked with 2×2cm squares (figure 1a). Under sterile, laminar air flow, a 25uL volume of *M. ulcerans* culture in different dilutions was spotted on the pigskin and left to air dry for approximately 5 minutes, after which the surfaces were swabbed (see below). A 7H9 sterile media control (blank) was also included (figure 1b). For each 2×2cm sample area, a sterile swab dipped once in saline was used. The swab was held like a pencil and swabbed firmly twice in both horizontal and vertical directions (figure 1c), to align with instructions provided to volunteers in the information packs for human skin swab collection. Each swab was placed in a 15mL Falcon tube and stored at 4°C until testing.

**Figure 1:**
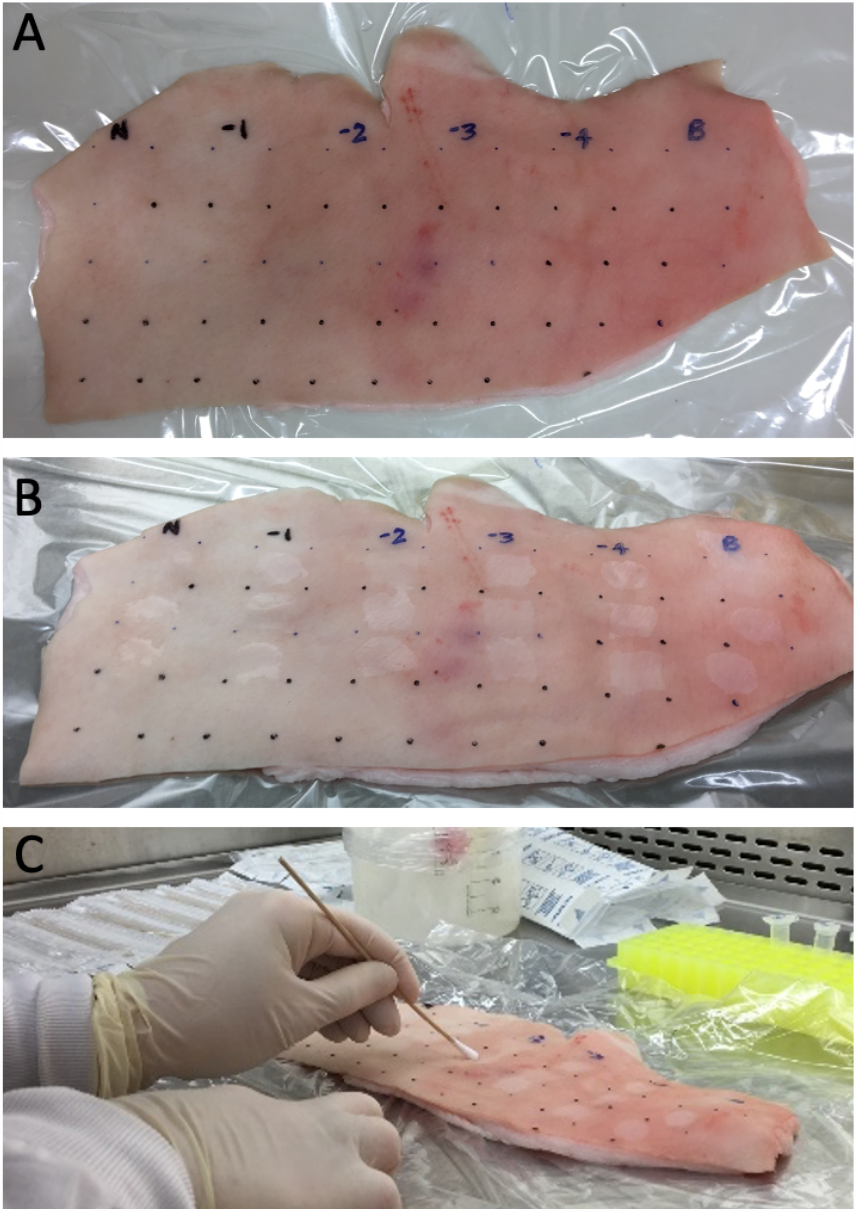
Preparation, *M. ulcerans* application and swabbing of fresh pigskin from part of one experiment. (A) The pigskin was divided in 2×2cm squares using a ruler and marker pen; (B) *M. ulcerans* culture and culture dilutions were applied to each square and allowed to dry before swabbing. Note that N = neat culture, −1,-2, −3, −4 etc are the 10-fold dilutions, B = sterile 7H9 media negative control; (C) Demonstration of swabbing technique for recovery of *M. ulcerans* from the pigskin surface.

### Human volunteers

Volunteers were friends and family of patients attending a Buruli clinic at Austin Health in Melbourne. Former patients were contacted via their treating clinician and asked to approach their friends and family. Adults >18 years who were willing to participate and spend >4 hours outside in a BU endemic area on the day of swab collection were included. Control participants were contacted directly through university networks and were included if they had not visited a known endemic area in the previous 6 months, were >18 years and would spend >4 hours outside in a non-endemic area in or around Melbourne on the day they collected their swabs. Recruitment was conducted from November 2018 until April 2019, corresponding to late spring, summer and early autumn, the expected peak transmission period.

Participating volunteers collected their own swabs after at least 4 hours (consecutive or non-consecutive) outdoors in their endemic or control environment. They were asked not to wash until the swab collection process was completed. Participants received a research kit containing a cover letter, four skin swabs in containers, sterile saline, a participant information/consent form, a questionnaire, skin swab instruction, a pre-addressed return bubble-wrap envelope and research recruitment flyers to pass on to additional friends and family.

Participants were asked to swab four areas with two or three separate swabs: back of elbows, back of wrists/forearms, back of calves and the ankles. One swab was to be used for both sides, except when the participant had a known BU, then that side of the body was to be avoided completely. These sites were chosen based a previous study which determined that Buruli occurs most frequently at these body locations [19].

### Ethical approvals

The study was approved by the Austin Health research ethics committee under reference number HREC/17/Austin/58. This observational clinical study was registered with Australia and New Zealand Clinical Trial Registry (ACTRN12619000998145).

## Results

### Validation of swab assay using pigskin

To assess the validity of swabbing skin to recover *M. ulcerans* for subsequent IS2404 qPCR detection, we established a model assay using a dilution series of *M. ulcerans* applied to sections of fresh pigskin. Two, 10-fold dilution series from a 10mL stationary phase culture of *M. ulcerans* (strain JKD8049) were prepared from 10^−1^ to 10^−6^. These two replicate series were labelled ‘A’ and ‘B’ and had undiluted (neat) bacterial concentrations of 9.3 × 10^5^ CFU/mL and 1.0 × 10^6^ CFU/mL respectively, as estimated by spot plating the dilutions (see methods), and represented an average *M. ulcerans* concentration of 5500 CFU/cm^2^. A 25uL aliquot of the neat culture and the six dilutions for both ‘A’ and ‘B’ preparations were spotted onto the pigskin (see example layout, figure 1B). The areas were swabbed (see methods), DNA was extracted and IS2404 qPCR performed. Concentrations of *M. ulcerans* on pigskin down to 0.06 CFU/cm^2^ were detected for both replicates ‘A’ and ‘B’. There was a premature flattening of the linear curve for the qPCR assay that began below concentrations of 10 CFU/cm^2^, from which a limit-of-detection for this assay was set at ≤ Ct 35 (highest Ct value of any replicate was 34.34) (Figure 2, Table S1). Unexposed control areas of pigskin (where Middlebrook 7H9 media only was applied) tested negative by IS2404 qPCR.

**Figure 2:**
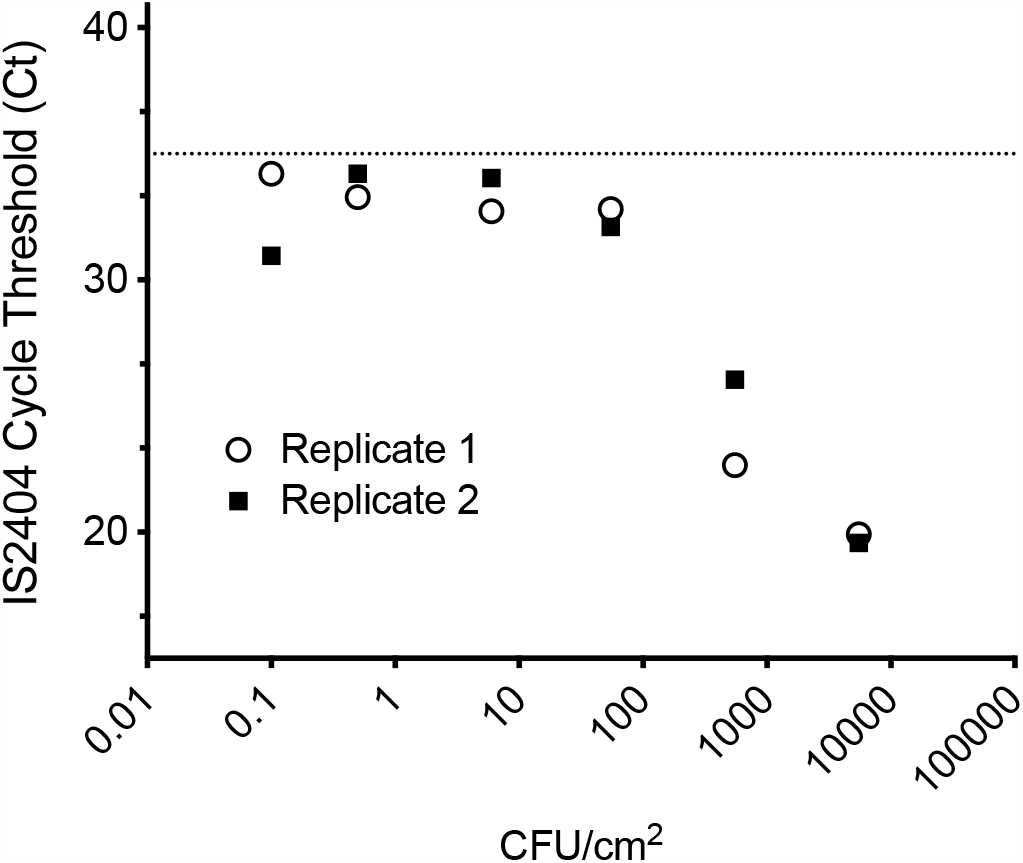
Limit-of-detection for IS2404 qPCR from swabbed pigskin. Plot showing Ct values for two independent *M. ulcerans* 10-fold dilution series used to inoculate fresh pigskin. The dotted line represents the limit-of-detection (Ct 35). Each dilution was tested by IS2404 qPCR in technical triplicates. Shown are the mean Ct values and standard deviation for the technical replicates. Note that error bars are present but too small to resolve (refer Table S1).

### Screening of swab specimens from human skin

Self-collected skin swabs were returned by 37 people who had been present in known Buruli ulcer endemic areas as recorded on their study survey and 10 who recorded no contact with an endemic area either on the day of collection or for the preceding 6 months as per the study protocol definition of “control area”. The cohort of 47 consisted of 18 males and 29 females, all adults of various ages. Of the 37 participants from endemic areas, 8 had been diagnosed and treated for Buruli ulcer although did not have active lesions at the time of the study. Each participant returned 4 sets of swabs (combined left and right elbows, calves, wrists, ankles). We also tested 16 sets of 4 swabs from 10 “Researchers” who were members of the *Beating Buruli* research team [23] performing outdoor environmental research in Buruli endemic areas during period of this study (duties included setting mosquito traps and conducting possum excreta surveys).

Control, researcher and participant swabs were coded and mixed so their provenance was not apparent to our laboratory technician and were intermingled with DNA extraction blanks made by dipping sterile swabs in saline in a ratio of one blank for every 10 test samples. All swabs were then screened by IS2404 qPCR. All results from the human volunteers in the endemic and control zones and the 10 researchers were negative (Cts ≥ 35) (Table S2). This included all swabs from the 8 volunteers with recent treated Buruli.

We did identify three individuals who had high but consistent results in the CT range 37-39 in replicates. One sample was from an ankle swab from a male from an endemic area (2 replicates – Ct values 37.4,37.84). A second was from the elbow of a female from an endemic area (38.33, 38.49). The third was an ankle swab from one of the researchers (KV) who worked outdoors while collecting possum excreta, a significant proportion of which was later shown to be heavily contaminated with *M. ulcerans* [24]. Ct values from KV’s ankle swab in January 2019 were 37.11, 37.16 and 37.49. KV agreed to re-test himself twice more in February 2019 and March 2019 while working on the same project but did not again return consistent high-positive results. We also observed a scattering of other inconsistent results in the Ct range 37-39 (one replicate detected, others not detected) including one of our control blanks (Ct 38.4). There were also 7 subjects and multiple blanks with high positive results Ct = 40.

## Discussion

Buruli ulcer have become an important public health issue in Victoria, Australia [10]. In our temperate zone Buruli was once associated only with rural coastal areas with low density populations but has now also become established in suburban areas close to the centres of Melbourne and Geelong, where the populations at risk are much larger. While there has been significant progress with earlier diagnosis and effective simpler treatment [18, 25], no organised attempts to prevent Buruli ulcer have been possible because we lacked knowledge about how exactly *M. ulcerans* is being transmitted. In recent years we have systematically investigated the role of mosquitoes [13, 26, 27] as potential vectors and have established that possums (tree dwelling marsupials) are the major local environmental reservoir in our endemic regions [11]. The purpose of this new study was to investigate whether human skin contamination is a common first step in the acquisition of Buruli ulcer in people exposed to environmental contamination which could then be followed by a range of chance inoculation events including but not restricted to mosquito bites. The answer to this question is needed to guide public health interventions that can prevent beyond just emphasising early diagnosis.

Using a fresh pig-skin model we validated a skin swab assay and used it to screen adult volunteers exposed outside during peak Buruli transmission season. All results were negative within the established range of the assay. Among these, these were three individuals who had consistent high-positive results (replicates positive) with Ct values above the defined limit-of-detection. One individual agreed to be retested twice more after re-exposure but returned negative results. We think the best interpretation of all these findings is that they were non-specific false positives.

Our study has some limitations. While 47 individuals with exposure to endemic areas participated and returned negative results, the 95% population confidence intervals around our point estimate of 0% is 0-8%, so a larger study may have identified a lower rate of skin contamination that we were not able to detect. Secondly, it is possible that a different method of *M. ulcerans* recovery from skin using detergent or soap (for example) may have been more sensitive than the saline swab method we employed. Against this, our limit-of-detection of 0.06 CFU/cm^2^ is indicative of a very sensitive assay. One could argue that detection sensitivity beyond this threshold would be of limited significance with respect to disease transmission potential.

In conclusion, we found no evidence that adult humans with outdoor exposure in Buruli endemic areas of Victoria during peak transmission season have detectable skin contamination with *M. ulcerans*.

## Acknowledgements

We thank the volunteers in this study who generously agreed to participate and to recruit friends and relatives to assist us.

---

**Supplementary table 1 (S1).** All results from pigskin validation model. **Validation qPCR results**

| Table S1: Validation qPCR results |  |  |  |
| --- | --- | --- | --- |
| Well | Sample | Cp FAM IS2404 | Cp VIC IPC |
| A1 | 10-1 A | 20.03 | 26.83 |
| B1 | 10-1 A | 19.77 | 26.93 |
| C1 | 10-1 A | 19.96 | 26.94 |
| D1 | 10-1 B | 19.76 | 26.86 |
| E1 | 10-1 B | 19.46 | 26.86 |
| F1 | 10-1 B | 19.49 | 26.94 |
| A2 | 10-2 A | 22.78 | 26.9 |
| B2 | 10-2 A | 22.59 | 26.87 |
| C2 | 10-2 A | 22.59 | 26.89 |
| D2 | 10-2 B | 26.09 | 26.91 |
| E2 | 10-2 B | 26.02 | 27 |
| F2 | 10-2 B | 26.04 | 26.98 |
| A3 | 10-3 A | 32.83 | 26.92 |
| B3 | 10-3 A | 32.83 | 27.03 |
| C3 | 10-3 A | 32.76 | 27.03 |
| D3 | 10-3 B | 32.34 | 27.05 |
| E3 | 10-3 B | 32.15 | 27.04 |
| F3 | 10-3 B | 31.82 | 27.04 |
| A4 | 10-4 A | 32.7 | 26.92 |
| B4 | 10-4 A | 32.75 | 26.93 |
| C4 | 10-4 A | 32.71 | 27.01 |
| D4 | 10-4 B | 34.15 | 26.93 |
| E4 | 10-4 B | 34.29 | 26.96 |
| F4 | 10-4 B | 33.68 | 27.02 |
| A5 | 10-5 A | 33.39 | 26.89 |
| B5 | 10-5 A | 33.23 | 26.97 |
| C5 | 10-5 A | 33.23 | 26.97 |
| D5 | 10-5 B | 34.33 | 27.11 |
| E5 | 10-5 B | 34.18 | 27.1 |
| F5 | 10-5 B | 34.11 | 26.99 |
| A6 | 10-6 A | 34.34 | 26.93 |
| B6 | 10-6 A | 34.06 | 26.93 |
| C6 | 10-6 A | 34.23 | 26.92 |
| D6 | 10-6 B | 30.93 | 26.97 |
| E6 | 10-6 B | 31.01 | 27.03 |
| F6 | 10-6 B | 30.91 | 26.93 |
| A7 | 7H9 alone A | nd | 26.98 |
| B7 | 7H9 alone A | nd | 27 |
| C7 | 7H9 alone A | nd | 27.06 |
| D7 | 7H9 alone B | nd | 26.99 |
| E7 | 7H9 alone B | nd | 26.94 |
| F7 | 7H9 alone B | nd | 27.02 |
| A8 | Pig Skin alone A | nd | 26.87 |
| B8 | Pig Skin alone A | nd | 26.95 |
| C8 | Pig Skin alone A | nd | 26.96 |
| D8 | Pig Skin alone B | nd | 27.03 |
| E8 | Pig Skin alone B | nd | 27 |
| F8 | Pig Skin alone B | nd | 26.96 |
| A9 | human skin A | nd | 26.98 |
| B9 | human skin A | nd | 26.94 |
| C9 | human skin A | nd | 26.91 |
| D9 | blank-extraction-control | nd | 26.93 |
| E9 | blank-extraction-control | nd | 27.12 |
| F9 | blank-extraction-control | nd | 26.96 |
| A10 | human skin B | nd | 26.86 |
| B10 | human skin B | nd | 26.86 |
| C10 | human skin B | nd | 26.9 |
| D10 | blank-extraction-control | nd | 26.98 |
| E10 | blank-extraction-control | nd | 27.09 |
| F10 | blank-extraction-control | nd | 27.05 |
| A11 | no-template-control | nd | 26.8 |
| B11 | no-template-control | nd | 26.81 |
| C11 | no-template-control | nd | 26.81 |
| A12 | positive control Mu gDNA | 12.28 | 26.72 |
| B12 | positive control Mu gDNA | 12.43 | 26.82 |
| C12 | positive control Mu gDNA | 12.44 | 26.81 |

**Supplementary table 2 (S2).**
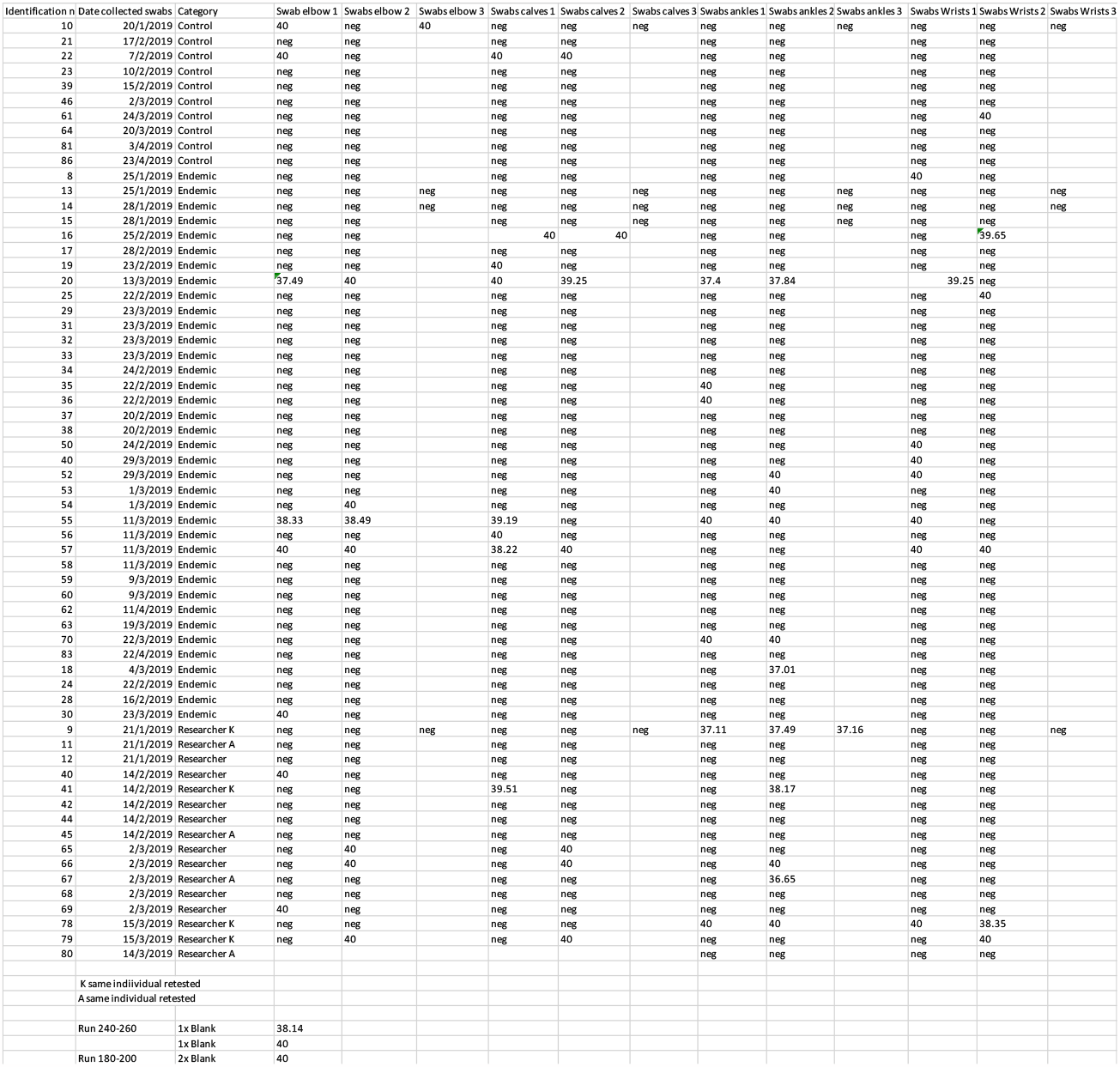
All results from human swabs. Blank cells indicate “not tested” as while all swabs were tested in duplicate at least, some were tested in triplicate.

## Notes

### Competing Interest Statement

The authors have declared no competing interest.

## References

1. Johnson PD, Stinear T, Small PL, Pluschke G, Merritt RW, Portaels F, et al. Buruli ulcer (M. ulcerans infection): new insights, new hope for disease control. PLoS medicine. 2005;2(4):e108. Epub 2005/04/21. doi: 10.1371/journal.pmed.0020108. PubMed PMID: 15839744; PubMed Central PMCID: PMCPMC1087202.

2. Friedman ND, Athan E, Walton AL, O’Brien DP. Increasing Experience with Primary Oral Medical Therapy for Mycobacterium ulcerans Disease in an Australian Cohort. Antimicrobial agents and chemotherapy. 2016;60(5):2692–5. Epub 2016/02/18. doi: 10.1128/aac.02853-15. PubMed PMID: 26883709; PubMed Central PMCID: PMCPMC4862461.

3. Phillips RO, Sarfo FS, Abass MK, Abotsi J, Wilson T, Forson M, et al. Clinical and bacteriological efficacy of rifampin-streptomycin combination for two weeks followed by rifampin and clarithromycin for six weeks for treatment of Mycobacterium ulcerans disease. Antimicrobial agents and chemotherapy. 2014;58(2):1161–6. Epub 2013/12/11. doi: 10.1128/aac.02165-13. PubMed PMID: 24323473; PubMed Central PMCID: PMCPMC3910847.

4. Friedman ND, Athan E, Hughes AJ, Khajehnoori M, McDonald A, Callan P, et al. Mycobacterium ulcerans disease: experience with primary oral medical therapy in an Australian cohort. PLoS neglected tropical diseases. 2013;7(7):e2315. Epub 2013/07/23. doi: 10.1371/journal.pntd.0002315. PubMed PMID: 23875050; PubMed Central PMCID: PMCPMC3715400.

5. Chauty A, Ardant MF, Marsollier L, Pluschke G, Landier J, Adeye A, et al. Oral treatment for Mycobacterium ulcerans infection: results from a pilot study in Benin. Clinical infectious diseases : an official publication of the Infectious Diseases Society of America. 2011;52(1):94–6. Epub 2010/12/15. doi: 10.1093/cid/ciq072. PubMed PMID: 21148526.

6. Nienhuis WA, Stienstra Y, Thompson WA, Awuah PC, Abass KM, Tuah W, et al. Antimicrobial treatment for early, limited Mycobacterium ulcerans infection: a randomised controlled trial. Lancet (London, England). 2010;375(9715):664–72. Epub 2010/02/09. doi: 10.1016/s0140-6736(09)61962-0. PubMed PMID: 20137805.

7. Phillips RO, Robert J, Abass KM, Thompson W, Sarfo FS, Wilson T, et al. Rifampicin and clarithromycin (extended release) versus rifampicin and streptomycin for limited Buruli ulcer lesions: a randomised, open-label, non-inferiority phase 3 trial. Lancet (London, England). 2020;395(10232):1259–67. Epub 20200312. doi: 10.1016/S0140-6736(20)30047-7. PubMed PMID: 32171422; PubMed Central PMCID: PMCPMC7181188.

8. Omansen TF, Erbowor-Becksen A, Yotsu R, van der Werf TS, Tiendrebeogo A, Grout L, et al. Global Epidemiology of Buruli Ulcer, 2010-2017, and Analysis of 2014 WHO Programmatic Targets. Emerging infectious diseases. 2019;25(12):2183–90. Epub 2019/11/20. doi: 10.3201/eid2512.190427. PubMed PMID: 31742506; PubMed Central PMCID: PMCPMC6874257.

9. Steffen CM, Freeborn H. Mycobacterium ulcerans in the Daintree 2009-2015 and the mini-epidemic of 2011. ANZ journal of surgery. 2018;88(4):E289–e93. Epub 2016/11/03. doi: 10.1111/ans.13817. PubMed PMID: 27804194.

10. Loftus MJ, Tay EL, Globan M, Lavender CJ, Crouch SR, Johnson PDR, et al. Epidemiology of Buruli Ulcer Infections, Victoria, Australia, 2011-2016. Emerging infectious diseases. 2018;24(11):1988–97. Epub 2018/10/20. doi: 10.3201/eid2411.171593. PubMed PMID: 30334704; PubMed Central PMCID: PMCPMC6199991.

11. Fyfe JA, Lavender CJ, Handasyde KA, Legione AR, O’Brien CR, Stinear TP, et al. A major role for mammals in the ecology of Mycobacterium ulcerans. PLoS neglected tropical diseases. 2010;4(8):e791. Epub 2010/08/14. doi: 10.1371/journal.pntd.0000791. PubMed PMID: 20706592; PubMed Central PMCID: PMCPMC2919402.

12. Carson C, Lavender CJ, Handasyde KA, O’Brien CR, Hewitt N, Johnson PD, et al. Potential wildlife sentinels for monitoring the endemic spread of human buruli ulcer in South-East australia. PLoS neglected tropical diseases. 2014;8(1):e2668. Epub 2014/02/06. doi: 10.1371/journal.pntd.0002668. PubMed PMID: 24498452; PubMed Central PMCID: PMCPMC3907424.

13. Johnson PD, Azuolas J, Lavender CJ, Wishart E, Stinear TP, Hayman JA, et al. Mycobacterium ulcerans in mosquitoes captured during outbreak of Buruli ulcer, southeastern Australia. Emerging infectious diseases. 2007;13(11):1653–60. Epub 2008/01/26. doi: 10.3201/eid1311.061369. PubMed PMID: 18217547; PubMed Central PMCID: PMCPMC3375796.

14. Wallace JR, Mangas KM, Porter JL, Marcsisin R, Pidot SJ, Howden B, et al. Mycobacterium ulcerans low infectious dose and mechanical transmission support insect bites and puncturing injuries in the spread of Buruli ulcer. PLoS neglected tropical diseases. 2017;11(4):e0005553. Epub 2017/04/15. doi: 10.1371/journal.pntd.0005553. PubMed PMID: 28410412; PubMed Central PMCID: PMCPMC5406025.

15. Williamson HR, Mosi L, Donnell R, Aqqad M, Merritt RW, Small PL. Mycobacterium ulcerans fails to infect through skin abrasions in a guinea pig infection model: implications for transmission. PLoS neglected tropical diseases. 2014;8(4):e2770. Epub 2014/04/12. doi: 10.1371/journal.pntd.0002770. PubMed PMID: 24722416; PubMed Central PMCID: PMCPMC3983084.

16. Loftus MJ, Trubiano JA, Tay EL, Lavender CJ, Globan M, Fyfe JAM, et al. The incubation period of Buruli ulcer (Mycobacterium ulcerans infection) in Victoria, Australia - Remains similar despite changing geographic distribution of disease. PLoS neglected tropical diseases. 2018;12(3):e0006323. Epub 2018/03/20. doi: 10.1371/journal.pntd.0006323. PubMed PMID: 29554096; PubMed Central PMCID: PMCPMC5875870.

17. Trubiano JA, Lavender CJ, Fyfe JA, Bittmann S, Johnson PD. The incubation period of Buruli ulcer (Mycobacterium ulcerans infection). PLoS neglected tropical diseases. 2013;7(10):e2463. Epub 2013/10/08. doi: 10.1371/journal.pntd.0002463. PubMed PMID: 24098820; PubMed Central PMCID: PMCPMC3789762.

18. Coutts SP, Lau CL, Field EJ, Loftus MJ, Tay EL. Delays in Patient Presentation and Diagnosis for Buruli Ulcer (Mycobacterium ulcerans Infection) in Victoria, Australia, 2011-2017. Tropical medicine and infectious disease. 2019;4(3). Epub 2019/07/07. doi: 10.3390/tropicalmed4030100. PubMed PMID: 31277453; PubMed Central PMCID: PMCPMC6789443.

19. Yerramilli A, Tay EL, Stewardson AJ, Kelley PG, Bishop E, Jenkin GA, et al. The location of Australian Buruli ulcer lesions-Implications for unravelling disease transmission. PLoS neglected tropical diseases. 2017;11(8):e0005800. Epub 2017/08/19. doi: 10.1371/journal.pntd.0005800. PubMed PMID: 28821017; PubMed Central PMCID: PMCPMC5584971.

20. Sexton-Oates NK, Stewardson AJ, Yerramilli A, Johnson PDR. Does skin surface temperature variation account for Buruli ulcer lesion distribution? PLoS neglected tropical diseases. 2020;14(4):e0007732. Epub 20200420. doi: 10.1371/journal.pntd.0007732. PubMed PMID: 32310955; PubMed Central PMCID: PMCPMC7192506.

21. Tobias NJ, Seemann T, Pidot SJ, Porter JL, Marsollier L, Marion E, et al. Mycolactone gene expression is controlled by strong SigA-like promoters with utility in studies of Mycobacterium ulcerans and buruli ulcer. PLoS neglected tropical diseases. 2009;3(11):e553. Epub 2009/11/26. doi: 10.1371/journal.pntd.0000553. PubMed PMID: 19936295; PubMed Central PMCID: PMCPMC2775157.

22. Fyfe JA, Lavender CJ, Johnson PD, Globan M, Sievers A, Azuolas J, et al. Development and application of two multiplex real-time PCR assays for the detection of Mycobacterium ulcerans in clinical and environmental samples. Applied and environmental microbiology. 2007;73(15):4733–40. Epub 2007/05/29. doi: 10.1128/aem.02971-06. PubMed PMID: 17526786; PubMed Central PMCID: PMCPMC1951036.

23. Department of Health V. Beating Buruli in Victoria 2018 [cited 2023 18/1/2023]. Available from: https://www.health.vic.gov.au/infectious-diseases/beating-buruli-in-victoria.

24. Koen Vandelannoote, Andrew H. Buultjens, Jessica L. Porter, Anita Velink, John R. Wallace, Kim R. Blasdell, et al. Structured surveys of Australian native possum excreta predict Buruli ulcer occurrence in humans (currently in review). BioRx. 2023; doi: https://doi.org/10.1101/2022.11.16.516821.

25. O’Brien DP, Jenkin G, Buntine J, Steffen CM, McDonald A, Horne S, et al. Treatment and prevention of Mycobacterium ulcerans infection (Buruli ulcer) in Australia: guideline update. The Medical journal of Australia. 2014;200(5):267–70. doi: 10.5694/mja13.11331. PubMed PMID: 24641151.

26. Quek TY, Athan E, Henry MJ, Pasco JA, Redden-Hoare J, Hughes A, et al. Risk factors for Mycobacterium ulcerans infection, southeastern Australia. Emerging infectious diseases. 2007;13(11):1661–6. Epub 2008/01/26. doi: 10.3201/eid1311.061206. PubMed PMID: 18217548; PubMed Central PMCID: PMCPMC3375781.

27. Lavender CJ, Fyfe JA, Azuolas J, Brown K, Evans RN, Ray LR, et al. Risk of Buruli ulcer and detection of Mycobacterium ulcerans in mosquitoes in southeastern Australia. PLoS neglected tropical diseases. 2011;5(9):e1305. Epub 2011/09/29. doi: 10.1371/journal.pntd.0001305. PubMed PMID: 21949891; PubMed Central PMCID: PMCPMC3176747.

